# Acidic fibroblast growth factor underlies microenvironmental regulation of MYC in pancreatic cancer

**DOI:** 10.1101/709261

**Authors:** Sohinee Bhattacharyya, Chet Oon, Aayush Kothari, Wesley Horton, Jason Link, Rosalie C. Sears, Mara H. Sherman

## Abstract

Despite a critical role for MYC as an effector of oncogenic RAS, strategies to target MYC activity in RAS-driven cancers are lacking. Oncogenic KRAS is insufficient to drive tumorigenesis, while addition of modest overexpression of MYC drives robust tumor formation, suggesting that mechanisms beyond the RAS pathway play key roles in MYC regulation and RAS-driven tumorigenesis. Here we show that acidic fibroblast growth factor (FGF1) derived from cancer-associated fibroblasts (CAFs) cooperates with cancer cell-autonomous signals to increase MYC level, promoter occupancy, and activity. FGF1 is necessary and sufficient for paracrine regulation of MYC protein stability, signaling through AKT and GSK-3β. These signals cooperate with, but are distinct from, cell-autonomous signals from oncogenic KRAS which stabilize MYC. Human pancreatic cancer specimens reveal a strong correlation between stromal CAF content and MYC protein level in the neoplastic compartment, and identify CAFs as the specific source of FGF1 in the tumor microenvironment. Together, our findings demonstrate that MYC is coordinately regulated by cell-autonomous and microenvironmental signals, and establish CAF-derived FGF1 as a novel paracrine regulator of oncogenic transcription.

**Statement of significance:** Our work highlights an unanticipated role for the tumor microenvironment in the regulation of MYC protein stability in pancreatic cancer cells and identifies CAF-derived FGF1 as a novel, specific paracrine regulator of MYC.

## Introduction

The *KRAS* oncogene is mutated in >90% of pancreatic ductal adenocarcinoma (PDAC) (1) and oncogenic KRAS is critical for PDAC initiation and maintenance (2, 3), making KRAS and its key effectors appealing targets for therapy. The oncogenic transcription factor MYC is well established as a critical effector of oncogenic RAS in multiple tumor types (4-7). In genetically engineered mouse models of lung and pancreatic cancer (8, 9), oncogenic KRAS is insufficient to drive tumorigenesis, while addition of modest MYC overexpression from the *Rosa26* locus drives robust tumor formation (10, 11), suggesting that mechanisms beyond the RAS pathway play key roles in MYC regulation and RAS-driven tumorigenesis. We have previously found that stromal cues from PDAC CAFs induce a transcriptional program in PDAC cells that significantly overlaps with the transcriptional network regulated by oncogenic KRAS (2, 12). This overlap suggests a gene-regulatory point of convergence for cell-autonomous and microenvironmental signals. The KRAS-regulated network was previously attributed to MYC-dependent transcription (2), but a role for a fibroinflammatory tumor microenvironment in paracrine regulation of MYC has not been established. MYC protein is very short-lived, and its expression and activity are exquisitely dependent on mitogenic signals (13, 14). While KRAS-mutant PDAC cells exhibit MYC protein stabilization downstream of ERK1/2 (15) or ERK5 (16), we reasoned that oncogenic levels of MYC in vivo may result from additional signals from the tumor microenvironment, and specifically from stromal CAFs.

## Results

To address a role for CAFs in paracrine regulation of MYC, we applied conditioned media (CM) from primary human PDAC CAFs to PDAC cells, and assessed MYC level. Both Western blot and immunofluorescence microscopy demonstrated that the CAF secretome acted in a paracrine manner to increase MYC protein level (Fig. 1A-D, Supplementary Fig. 1A-D), peaking by 3h. Importantly, a non-cancer associated human pancreatic stellate cell (hPSC) line did not induce MYC under the same experimental conditions (Fig. 1A), suggesting specificity for CAFs and arguing against a non-specific effect of CM. These increases were more pronounced in the soluble than the insoluble nuclear fraction (at 400mM NaCl); as MYC is found in both fractions (17), we examined total nuclear extracts moving forward. Before performing mechanistic studies, we assessed the relationship between stromal CAF content and MYC level in human PDAC. Immunohistochemical analysis revealed a strong correlation between MYC protein level in keratin-positive PDAC cells and *α*-smooth muscle actin (αSMA)-positive CAF density among human PDAC samples (Fig. 1E,F), supporting the notion that CAFs may signal in a paracrine manner to augment MYC expression in the neoplastic compartment. Importantly, this was not a reflection of increased density of cancer cells among stroma-rich PDAC regions, as we saw no correlation between keratin and αSMA in these tissues (Supplementary Fig. 1E). We stained for MYC pS62 in these analyses as total MYC antibodies did not yield specific staining of our human PDAC tissues (see Materials and Methods). To begin to understand the mechanism by which CAFs increase MYC protein levels in PDAC cells, we tested MYC RNA and protein stability under control and CAF CM-treated conditions. CAF CM significantly increased MYC protein stability (Fig. 1G,H), while RNA stability was not significantly changed (Fig. 1I), though we noted that MYC RNA at steady state was increased with CM (Supplementary Fig. 2C). CAF CM also increased PDAC cell proliferation (Fig. 1J), as expected for conditions which augment MYC level and activity and consistent with previous results (18). These results suggest that PDAC CAFs signal to stabilize MYC in the neoplastic compartment in a paracrine manner, an interaction that is reflected in human PDAC samples.

**Figure 1:**
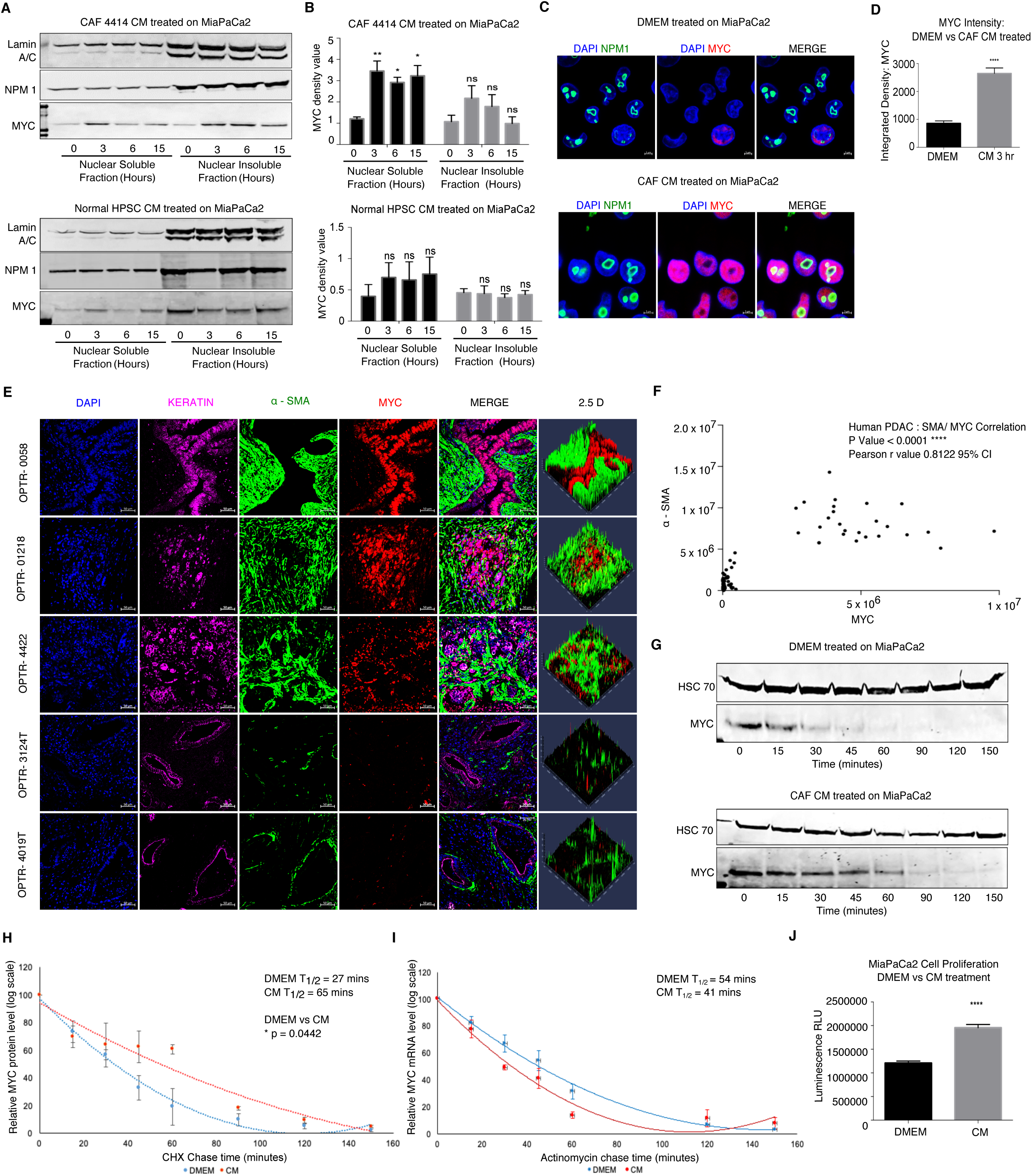
PDAC CAF-derived factors increase MYC level in PDAC cells. **A,** Western blots showing MYC levels in PDAC cells after treatment with CM from CAFs or hPSC for the indicated duration. Lamin A/C and NPM1 (Nucleophosmin) are loading controls. **B,** Quantification of MYC expression normalized to NPM1 in MiaPaCa2 cells treated with CAF 4414 or hPSC CM for the indicated duration. *p < 0.05, **p < 0.01 by two-way ANOVA (n = 3 biological replicates). **C,** Immunofluorescence (IF) microscopy showing MYC levels in PDAC cells treated with DMEM or CAF CM for 3h. NPM1 shows localization of nucleoli; DAPI stains nuclei. Scale bar = 5μm. **D,** Quantification of MYC intensity in **C**. ****p < 0.0001 by Student’s t-test (n = 3 biological replicates). 78 fields of view were quantified per treatment condition. **E,** Fluorescent immunostaining of human PDAC patient tissue samples. DAPI stains nuclei, pan-KRT (keratin) stains PDAC cells, *α*SMA stains CAFs. Scale bar = 50μm. **F,** Quantification of integrated density of MYC in KRT^+^ cells versus *α*SMA in 80 fields of view from n = 5 human PDAC specimens. **G,** MiaPaCa2 cells were treated with DMEM or CAF 4414 CM for 3h, then treated with cycloheximide and harvested at the indicated time points to measure MYC protein level by Western blot. HSC70 is a loading control. **H,** Quantification of MYC levels in **G** (n = 3 biological replicates). p-value reflects comparison of t_half_ from biological triplicates by Student’s t-test. **I,** MiaPaCa2 cells were treated as in **G**, then treated with Actinomycin D and harvested at the indicated time points for qRT-PCR analysis of MYC mRNA levels (n = 3 biological replicates). Data were normalized to ACTB as a housekeeping gene. For **H** and **I**, data were plotted on a semi-log scale and best-fit lines were calculated using linear regression. **J,** Proliferation assay on MiaPaCa2 cells treated with DMEM or CAF 4414 CM for 72h. ****p < 0.0001 by Student’s t-test (n = 3 biological replicates).

We next aimed to identify the specific CAF-derived factor that stabilizes MYC in PDAC cells. As our hPSC line did not induce MYC via secreted factors, we used this line as a basis for comparison. We extracted a list of secreted growth factors, cytokines, and chemokines expressed in PDAC CAFs from a published RNA-seq dataset (19), and compared expression of each factor in hPSC versus CAFs (Fig. 2A). This comparison identified 6 factors with at least 3-fold higher expression in CAFs than hPSC (FGF2 was included at 2.9-fold). We then tested neutralizing antibodies against each of these 6 candidate factors in CAF CM to determine whether inhibition of an individual factor could suppress paracrine induction of MYC. Though SHH was not detected in the RNA-seq dataset, it is variably expressed in primary CAFs including the CAF sample used for screening, so we included an antibody against SHH as well as NOTCH1 and NOTCH3 based on prior work linking them to MYC (20). Our top candidate from this neutralizing antibody screen was acidic fibroblast growth factor (FGF1) (Fig. 2B, Supplementary Fig. 2A). FGF1 has not previously been linked to MYC regulation, and has not been studied functionally in the pancreatic tumor microenvironment. We thus confirmed that primary PDAC CAFs secrete FGF1. ELISA assays revealed that PDAC CAFs secrete abundant FGF1, while expression was restricted to basal levels in CM from hPSC and from human PDAC cells (Fig. 2C), consistent with a role in paracrine signaling. We also examined FGF1 expression by immunofluorescence microscopy, which similarly showed detectable FGF1 in PDAC CAFs but not in hPSC or PDAC cells (Fig. 2D). FGF1 signals through the four-member family of fibroblast growth factor receptors (FGFRs); to assess whether FGFRs are expressed in PDAC cells, we stained for FGFR1, a family member recently implicated as a target for combination therapy in PDAC (21), and found that this receptor was expressed in PDAC cells; interestingly, this receptor was also highly expressed in hPSC but was found at a very low level in CAFs, showing a reciprocal staining pattern to that of FGF1 (Fig. 2D). Consistent with results in cultured cells, human PDAC tissues showed robust staining for FGFR1, while CAFs were mostly negative (Fig. 2E). We next assessed both the expression and the cellular source of FGF1 in human PDAC. As FGF1 is a secreted factor, instead of immunohistochemistry we probed FGF1 expression using RNA fluorescence in situ hybridization (RNA-FISH), and co-stained for ACTA2 (encoding αSMA). FGF1 signal was restricted to ACTA2-positive cells, though not all ACTA2-positive cells express FGF1 (Fig. 2F), suggesting that the *α*SMA-positive myofibroblastic CAF (myCAF) subset produces FGF1 in PDAC (22). These results suggest that CAFs are the principal cellular source of FGF1 in the PDAC microenvironment, and that the FGF1-FGFR axis is expressed in human PDAC.

**Figure 2:**
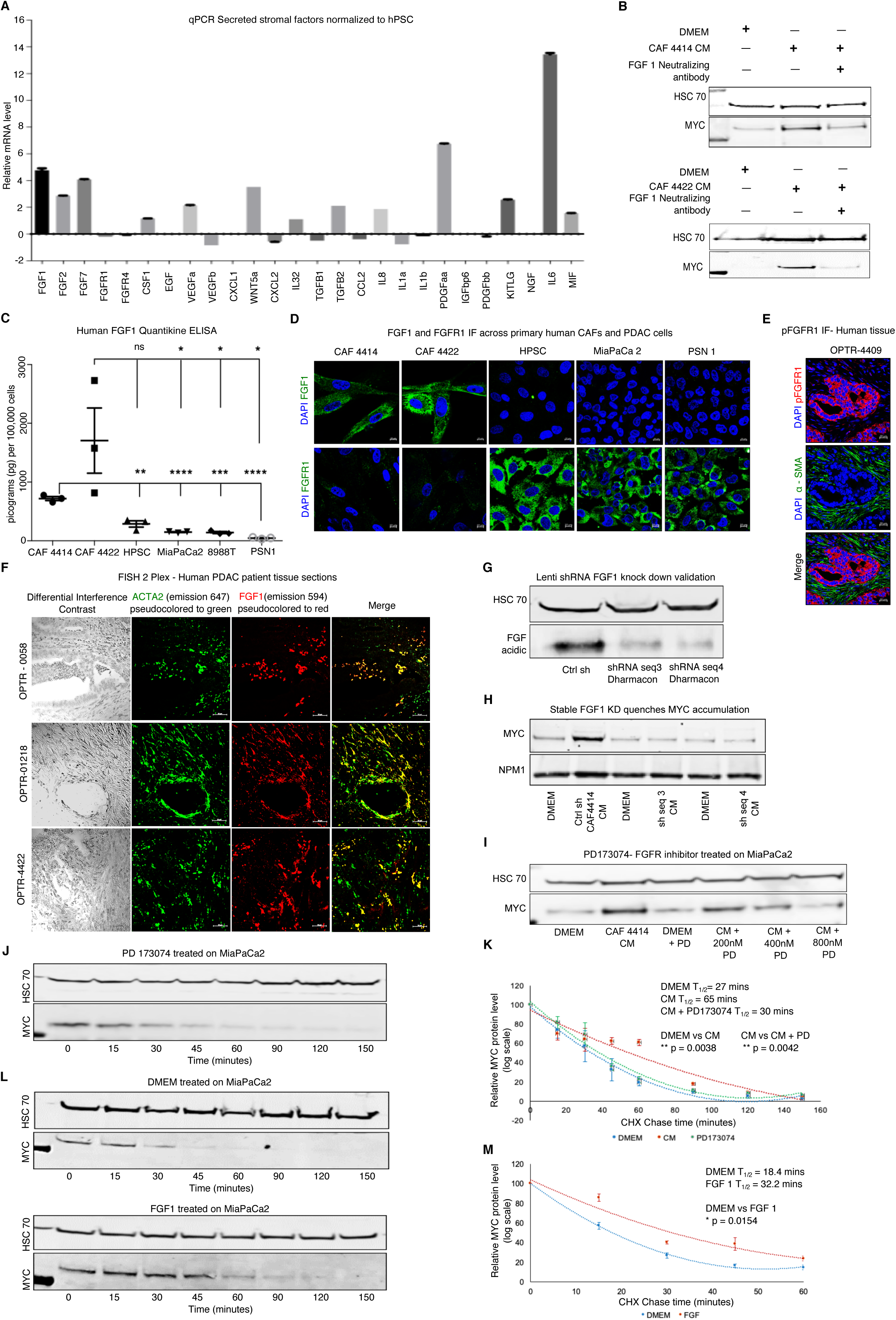
Stroma-derived FGF1 is necessary and sufficient for paracrine regulation of MYC. **A,** qRT-PCR for the indicated secreted factors in CAF 4414 and hPSC (n = 3 biological replicates). Data are plotted as fold change in CAF 4414 compared to hPSC (set to 1); all values were normalized to 36B4 as a housekeeping gene. **B,** CAF 4414 or 4422 CM was incubated alone or with FGF1 neutralizing antibody for 1h at room temperature, then added to MiaPaCa2 cells for 3h. MYC levels in nuclear extracts were analyzed by Western blot (representative of 3 biological replicates). **C,** ELISA for FGF1 in CM from the indicated cell lines. Data presented as mean ± SEM, **p < 0.0025, *** p < 0.0001, ****p < 0.0001 by Student’s t-test (n = 3 biological replicates). **D,** Immunofluorescence staining for FGF1 or FGFR1 in the indicated cell lines. Scale bar = 10μm. **E,** Fluorescent immunostaining for pFGFR1 in human PDAC samples (representative of n = 3 patient samples). Scale bar = 50μm. **F,** RNA fluorescent in situ hybridization for ACTA2 (encoding *α*SMA) and FGF1 in human PDAC (representative of n = 6 patient samples). Scale bar = 50μm. Differential interference contrast (left column) shows tissue context. **G,** Western blots showing FGF1 knockdown in CAF 4414 knockdown lines (scramble control versus 2 distinct shFGF1 sequences). **H,** Western blot for MYC in MiaPaCa2 cells after treatment with DMEM or CM from the indicated CAF 4414 line for 3h (representative of n = 3 biological replicates). **I,** Western blot for MYC in MiaPaCa2 cells after treatment with DMEM or CM +/-FGFR inhibitor PD173074 at the indicated concentrations for 3h (representative of n = 3 biological replicates). **J,** MiaPaCa2 cells were treated with DMEM or CAF 4414 CM for 3h, then treated with cycloheximide +/-800nM PD173074 for the indicated duration and MYC levels were analyzed by Western blot. **K,** Quantification of MYC levels in **J** (n = 3 biological replicates). p-value reflects comparison of t_half_ from biological triplicates by Student’s t-test. **L,** MiaPaCa2 cells were treated with DMEM alone or supplemented with 50 pg/ml human FGF1 for 3h, then treated with cycloheximide for the indicated duration and MYC levels were analyzed by Western blot. **M,** Quantification of MYC levels in **l** (n = 3 biological replicates). p-value reflects comparison of t_half_ from biological triplicates by Student’s t-test.

We next performed functional studies to assess the significance of FGF1 in paracrine regulation of MYC. To this end, we stably knocked down FGF1 in primary PDAC CAFs using two independent hairpin sequences (Fig. 2G). CM from these knockdown lines resulted in markedly reduced MYC induction in PDAC cells compared to controls (Fig. 2H). To further examine this functional connection with pharmacologic inhibitors, and to probe a role for FGFRs in MYC regulation, we incubated PDAC cells in CAF CM in the presence or absence of FGFR inhibitor PD173074 or Debio-1347. Both inhibitors blocked induction of MYC by CAF-derived signals (Fig. 2I, Supplementary Fig. 2B). FGFR inhibitor prevented the stroma-inducible increase in MYC half-life, such that CM containing FGFR inhibitor yielded MYC half-life measurements similar to untreated controls (Fig. 2J,K). While these results together suggested that FGF1 is necessary to stabilize MYC, we next tested whether this factor is sufficient for MYC stabilization. Using the lowest concentration of FGF1 found in CAF CM in our ELISA assays (Fig. 2C), we found that recombinant human FGF1 was sufficient to increase MYC protein in PDAC cells and significantly increased MYC stability (Fig. 2L,M). FGF1 alone did not induce MYC RNA, and FGFR inhibition did not suppress the induction of MYC mRNA by CAF CM (Supplementary Fig. 2C,D), suggesting that MYC protein stabilization is the relevant mechanism of MYC induction by stroma-derived FGF1. Together, these results suggest that CAF-derived FGF1 is necessary and sufficient for paracrine stabilization of MYC protein in PDAC cells.

We next queried the pathway downstream of FGF1-FGFR signaling that results in stabilization of MYC. In RAS-mutant cancer cells, MYC is classically stabilized by phosphorylation on S62 downstream of the MAPK pathway; further phosphorylation on T58 targets MYC for S62 dephosphorylation and subsequent proteasomal degradation (23). AKT activation can result in an inhibitory phosphorylation event on GSK-3β, preventing phosphorylation of MYC T58 by GSK-3β and further promoting MYC protein stability (24). We found that CAF CM induced phosphorylation of FGFR1, as well as downstream AKT and GSK-3β (Fig. 3A, Supplementary Fig. 3A,B). ERK phosphorylation was high but unchanged by CM at this time point consistent with previous studies (25), suggesting that MYC stabilization by the stroma is distinct from cell-autonomous RAS-MAPK signaling. Total MYC and pS62 MYC were increased to a similar extent by CAF CM, while pT58 MYC was induced to a lesser extent (Fig. 3B), raising the possibility that CAF CM suppresses MYC T58 phosphorylation. The relevant concentration of recombinant FGF1 was sufficient to induce phosphorylation of AKT and GSK-3β at time points consistent with MYC stabilization (Fig. 3C). Addition of FGFR inhibitor to CM blunted paracrine induction of AKT and GSK-3β phosphorylation and increased pT58 MYC levels relative to total MYC, while ERK phosphorylation was increased (Fig. 3D,E). To determine whether FGF1 is necessary for CAFs to activate this axis in PDAC cells, we looked at signaling after incubating PDAC cells with CM from our FGF1 knockdown CAFs. Surprisingly, paracrine induction of AKT and GSK-3β phosphorylation was suppressed in the absence of FGF1 (Fig. 3F). Though CAFs secrete numerous growth factors which can presumably activate AKT, these results suggest that stromal FGF1 creates a permissive context needed for robust paracrine regulation of AKT. To further test the role of AKT and GSK-3β in regulation of MYC by stromal signaling, we added an AKT inhibitor to CAF CM, which reduced GSK-3β phosphorylation and MYC induction (Fig. 3G,H). To provide genetic evidence for the roles of these kinases, we transduced PDAC cells with dominant-negative AKT, or with constitutively active GSK-3β. Both interventions reduced paracrine induction of MYC (Supplementary Fig. 3C-E). To further assess the significance of MYC T58 phosphorylation in paracrine regulation of MYC stability, we transduced PDAC cells with Flag-tagged WT or T58A MYC, then treated these cells with CAF CM. While WT MYC levels were increased by CM, T58A MYC was not affected (Fig. 3I), consistent with stromal suppression of T58 phosphorylation as a mechanism underlying the increase in MYC protein stability. Together, these results provide support for a central role for AKT and GSK-3β in FGF1-mediated stabilization of MYC.

**Figure 3:**
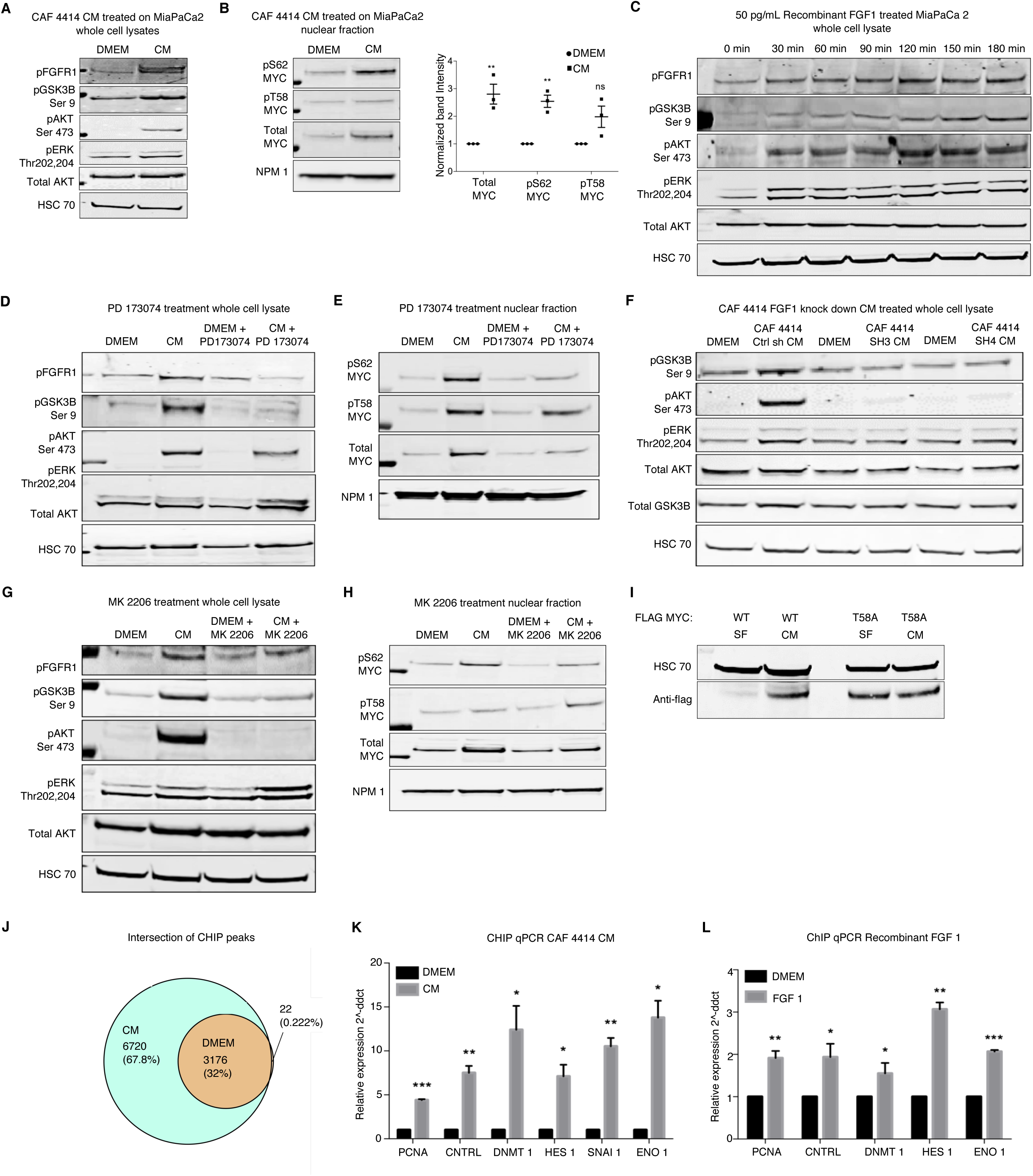
Paracrine FGF1 signaling augments MYC expression and stability via AKT/GSK3β signaling, and alters MYC chromatin occupancy. **A,** Western blots for the indicated signaling events after 3h treatment with DMEM or CAF 4414 CM. **B,** Western blots for the indicated MYC species after treatment as in **A**. Quantification appears to the right (n = 3 biological replicates), **p < 0.01 by Student’s t-test. **C,** Western blots for the indicated signaling events after a time course of treatment with 50 pg/ml recombinant human FGF1. **D,** Western blots for the indicated signaling events in MiaPaCa2 cells after 3h treatment with CAF 4414 CM +/-800nM PD173074. **E,** Western blots for the indicated MYC species in cells treated as in **D**. **F,** Western blots for the indicated signaling events in MiaPaCa2 cells treated with CM from control or FGF1 knockdown CAFs for 3h. **G,** Western blots for the indicated signaling events in MiaPaCa2 cells treated with CAF 4414 CM +/-AKT inhibitor MK2206 (10μM) for 3h. **H,** Western blots for the indicated MYC species from cells treated as in **G**. **I,** Western blots for Flag and HSC70 (loading control) in MIAPaCa2 cells transduced with WT or T58A Flag-tagged MYC and treated with CAF 4414 CM for 3h. **J,** Venn diagram showing overlap of significant MYC-bound peaks from ChIP-seq in MiaPaCa2 cells treated with DMEM or CAF 4414 CM for 3h (n = 2 biological replicates). **K,** ChIP-qPCR for MYC binding sites in the indicated differentially bound promoters, as determined by ChIP-seq, in MiaPaCa2 cells treated with DMEM or CAF 4414 CM for 3h (n = 2 biological replicates). **L,** ChIP-qPCR for MYC binding sites in the indicated differentially bound promoters in MIAPaCa2 cells treated with 50pg/ml FGF1 for 3h (n = 2 biological replicates). For **K** and **L**, *p < 0.05, **p < 0.01, ***p < 0.001 by Student’s t-test.

We next analyzed MYC genomic localization to better understand the consequences of increased MYC protein downstream of stromal FGF1. To this end, we performed ChIP-seq for MYC in PDAC cells incubated in control medium or CAF CM. As in previous studies featuring varying levels of MYC (26, 27), we find that nearly all MYC-bound sites under control conditions were also occupied under CM-stimulated conditions; CM resulted in 6,720 additional MYC binding sites (Fig. 3J). Unique MYC-bound promoters upon CM treatment were associated with genes involved in post-transcriptional regulatory mechanisms including RNA processing and ribosome biogenesis (Supplementary Fig. 4A). Further, commonly bound peaks generally had a higher ChIP-seq signal under CM-treated conditions (Supplementary Fig. 4B). Motif analysis showed that a consensus MYC binding site was the top enriched motif (p = 1 × 10^−100^) (Supplementary Fig. 4C), supporting the specificity of the immunoprecipitation for MYC. ChIP-qPCR for differentially bound loci validated our ChIP-seq results with CM from 2 independent CAF lines (Fig. 3K, Supplementary Fig. 4D), and these differentially bound genes also revealed significant changes in gene expression upon CM treatment (Supplementary Fig. 4E). Importantly, FGF1 was sufficient to increase MYC binding to these loci (Fig. 3L). These results demonstrate that stromal cues increase MYC promoter occupancy in PDAC cells, consistent with increased protein levels.

As MYC activity is principally associated with cellular growth control, we next assessed a role for the FGF1-FGFR axis in PDAC cell proliferation. While control CAF CM increased PDAC cell proliferation, this effect was significantly reduced with CM from FGF1 knockdown CAFs (Supplementary Fig. 5A,B). Similarly, FGFR inhibition significantly reduced stroma-inducible proliferation (Fig. 4A). To address the link between FGF1-FGFR signaling and MYC levels in vivo, we suppressed this signaling axis in three distinct models. First, we performed subcutaneous transplantation of human PDAC cells together with control or FGF1 knockdown CAFs, into the flanks of nude mice. Quantification of MYC intensity within KRT-positive cells showed that MYC levels in PDAC cells were significantly reduced when co-transplanted with FGF1-knockdown CAFs compared to controls (Fig. 4B,C). Next, we performed orthotopic transplantation of the murine KPC cell lines FC1199 and FC1245 into pancreata of immune-competent C57BL/6J hosts. Once tumors were detectable by ultrasound, mice were treated with vehicle or bioavailable FGFR inhibitor BGJ 398 for 9 days. FGFR inhibition resulted in a significant reduction in MYC protein level in PDAC cells in both models (Fig. 4D-G). While FGFR inhibition significantly suppressed tumor growth compared to controls (Fig. 4H), tumors did continue to grow. We hypothesized that inhibition of parallel MAPK signaling and FGF1-FGFR signaling would further reduce MYC levels by blocking 2 MYC-stabilizing pathways, and further reduce tumor growth. We first tested this hypothesis in vitro, and found that the combination of MEK inhibitor trametinib and BGJ 398 reduced paracrine induction of AKT and MYC compared to either single agent (Supplementary Fig. 5C). Combined treatment with trametinib and BGJ 398 significantly reduced tumor growth in vivo compared to either drug alone (Fig. 4H), consistent with a reduction in MYC levels in PDAC cells (Fig. 4F,G) and consistent with a previous study of trametinib and FGFR1 inhibition (21). Taken together, our results highlight a specific role for CAF-derived FGF1 as a paracrine regulator of MYC in pancreatic cancer.

**Figure 4:**
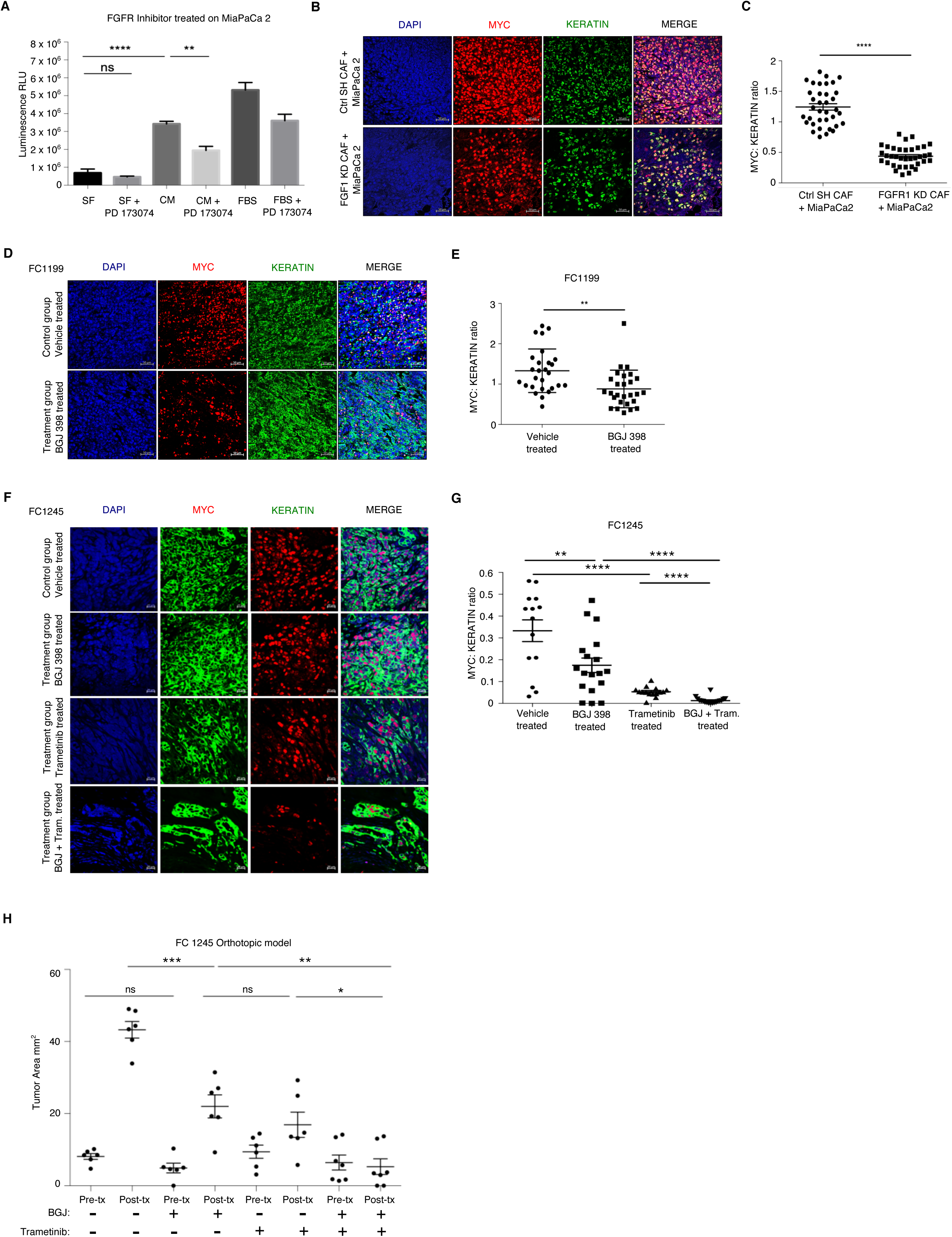
The FGF1/FGFR axis regulates MYC levels and PDAC growth in vivo. **A,** Proliferation assay for MiaPaCa2 cells treated with CAF 4414 CM +/-PD173074 (800nM) for 72h. **p < 0.01, ****p < 0.0001 by one-way ANOVA (n = 3 biological replicates). **B,** Fluorescent immunostaining of co-transplanted subcutaneous xenografts. Scale bar = 50μm. **C,** Quantification of samples in **B** to determine the ratio of MYC intensity within KRT^+^ cells over KRT signal. 36 fields of view were analyzed from n = 5 mice per condition. **D,** Fluorescent immunostaining of orthotopic transplants of FC1199 cells treated with vehicle or BGJ 398. **E,** Quantification of samples in **D** to determine the ratio of MYC intensity within KRT^+^ cells over KRT signal. 28 fields of view were analyzed from n = 3 mice per condition. For **C** and **E**, **p < 0.01, ****p < 0.0001 by Student’s t-test. **F,** Fluorescent immunostaining of orthotopic transplants of FC1245 cells from the indicated treatment groups. **G,** Quantification of samples in **F** to determine the ratio of MYC intensity within KRT^+^ cells over KRT signal (n = 3 mice per group, 6 fields imaged per mouse). **p < 0.01, ****p < 0.0001 by Student’s t-test. **H,** Tumor measurements by high-resolution ultrasound at the indicated time points in FC1245 orthotopic transplants (vehicle, BGJ 398, trametinib: n = 6; BGJ 398 + trametinib, n = 7). *p < 0.05, **p < 0.01, ***p < 0.001 by one-way ANOVA.

## Discussion

Though compelling prior studies have highlighted cell-autonomous mechanisms of MYC regulation in RAS-driven cancers (15, 16, 23, 24), here we show that MYC is further subject to regulation by soluble cues from stromal CAFs in pancreatic cancer. As fibroblasts evolved in part to facilitate wound repair, this may reflect a conserved mechanism by which mesenchymal cells signal to the epithelial compartment to promote regeneration and healing, as MYC has been previously implicated in the wound-healing process (28-30) as have MYC-regulated bioenergetics (31). While previous studies have demonstrated a critical role for MYC in regulation of the tumor microenvironment (10, 32), our findings suggest the existence of a feedback loop, in which cues from the fibroinflammatory microenvironment in turn augment MYC protein level and oncogenic activity. As PDAC rarely harbors PI-3K mutations (33), it was not entirely surprising that external signals can increase AKT phosphorylation in PDAC cells. However, we were surprised to find that FGF1 is required for paracrine AKT activation by CAFs, given the abundance of additional secreted factors from CAFs that may signal through growth factor receptors to activate AKT. These findings suggest that FGF1 creates a permissive context for activation of the AKT/GSK-3β axis, and future studies will aim to understand the broader proteomic network regulated by FGF1 in PDAC cells. Future efforts will also aim to discover additional therapeutic strategies which, together with FGF1/FGFR inhibition, improve outcome in RAS/MYC-driven cancers.

## Materials and Methods

### Animals

All experiments were reviewed and overseen by the institutional animal use and care committee at Oregon Health and Science University in accordance with NIH guidelines for the humane treatment of animals. C57BL/6J (000664) or NU/J (002019) mice from Jackson Laboratory were used for orthotopic transplant and xenograft experiments, respectively, at the ages denoted in the manuscript.

### Human tissue samples

Human patient PDAC tissue samples donated to the Oregon Pancreas Tissue Registry program (OPTR) in accordance with full ethical approval were kindly shared by Dr. Jason Link and Dr. Rosalie Sears.

### Cycloheximide chase

Cells were serum starved for 48 hours in serum free DMEM and following serum starvation, cells were either treated with DMEM or CAF derived CM for 3 hours following which, cells were treated with 50 μg/ml CHX for the indicated times. Then, the cells were lysed and whole cell lysates or nuclear lysates were prepared for Western blotting analysis. MYC levels relative to time point 0 are shown in all graphs of CHX experiments.

### Actinomycin D chase

Cells were serum starved for 48 hours in serum free DMEM and following serum starvation, cells were either treated with DMEM or CAF derived CM for 3 hours following which, Actinomycin D (5 µg/ml final concentration) was added and RNA was isolated at the times indicated. Analysis of MYC mRNA from PDAC cells was done by quantitative PCR using MYC specific primer pairs and threshold cycle (CT) values normalized using *ACTB* as a housekeeping gene. MYC mRNA levels relative to time point 0 are shown in all graphs of Actinomycin D chase experiments.

### Data Availability

All sequence data utilized for this study will be deposited in a publicly available database.

### Orthotopic Transplant/Allograft Model

The orthotopic transplant model used here was described previously (34). In brief, 8 week old wild-type male C57BL/6J mice were orthotopically transplanted as described previously with 10,000 FC1199 cells, isolated from PDAC in LSL-KrasG12D/+; Trp53R172H/+; Pdx1-Cre C57B6/J mice, obtained from the David Tuveson laboratory (Cold Spring Harbor Laboratory, NY), in 50% Matrigel. After ultrasound imaging to confirm the presence of PDAC, mice were randomized into treatment groups: mice that were treated by oral gavage daily with vehicle alone, with BGJ 398 (50mg/kg body weight), with trametinib (1mg/kg body weight), or BGJ 398 + trametinib. Tumors were measured and mice euthanized after 9 days of treatment, and pancreata were harvested, sliced, and flash frozen in liquid nitrogen or immediately fixed in formalin.

### Subcutaneous PDAC transplants

All surgical procedures and care of the animals were in accordance with institutional guidelines. A 100 μl volume of 1:1 mixture of Matrigel (354248; BD Biosciences) and 1 × 10^6^ MiaPaCa2 cells alone, or 1 × 10^6^ MiaPaCa2 cells plus 5 × 10^6^ control or shFGF1 CAFs was subcutaneously injected into NU/J (002019) mice flanks. Tumor size was measured every other day using digital calipers and tumor volume was estimated using the following formula: V = (LW2)/2, where V, volume (mm3); L, largest diameter (mm) and W, smallest diameter (mm). Mice were euthanized and tumors harvested on day 40 post-transplantation for analysis.

### Statistical Analysis

All statistical analyses were performed using GraphPad Prism 5.0 Software (Graph Pad Software Inc.).

## Acknowledgements

We thank the OHSU Massively Parallel Sequencing Shared Resource, Advanced Light Microscopy Shared Resource, and Histopathology Shared Resource for technical help; the OHSU Brenden-Colson Center for Pancreatic Care for providing primary PDAC reagents; Mark Berry and Cameron Roberts for technical support; and Sara Courtneidge and members of the Sherman lab for helpful discussion and feedback. This work was supported by funds from the National Cancer Institute (R00CA188259 and R01CA229580) and a Medical Research Foundation New Investigator Grant (to M.H.S.).

## Author Contributions

S.B. designed, performed, and analyzed all experiments with technical support from C.O. and A.K., and under the guidance of M.H.S. W.H. analyzed the ChIP-seq data. J.L. and R.C.S. assisted with generation of primary PDAC CAFs and tissue samples. S.B. and M.H.S. wrote the manuscript.

