## Supplement for "Acidic fibroblast growth factor underlies microenvironmental regulation of MYC in pancreatic cancer"

**Supplementary Figure Legends:**

**Supplementary Figure 1: PDAC CAF-derived factors increase MYC level in PDAC cells.**

**A,** Western blots showing MYC levels in Panc1 cells after treatment with CM from CAF4414 for the indicated duration. Lamin A/C, HSC70 and NPM1 (Nucleophosmin) are loading controls. **B,** Western blots showing MYC levels in PSN1 cells after treatment with CM from CAF4414 for the indicated duration. Lamin A/C, HSC70 and NPM1 (Nucleophosmin) are loading controls. **C,** Western blots showing MYC levels in MiaPaCa2 cells after treatment with CM from CAF4422 for the indicated duration. Lamin A/C, HSC70 and NPM1 (Nucleophosmin) are loading controls. **D,** Immunofluorescence (IF) microscopy showing MYC levels in 8988T PDAC cells treated with DMEM or CAF CM for 3h. NPM1 shows localization of nucleoli; DAPI stains nuclei. Scale bar = 5μm. **E,** Quantification of integrated density of Keratin versus αSMA in 50 fields across 5 different human PDAC patient samples.

**Supplementary Figure 2: Stroma-derived FGF1 is necessary and sufficient for paracrine regulation of MYC.**

**A,** CAF 4414 CM was incubated alone or with the indicated neutralizing antibodies for 1h at room temperature, then added to MiaPaCa2 cells for 3h. MYC levels in nuclear extracts were analyzed by Western blot (representative of 3 biological replicates). **B,** Western blot for MYC in MIAPaCa2 cells after treatment with DMEM or CM +/- FGFR inhibitor Debio 1347 at the indicated concentrations for 3h. **C,** qRT-PCR analysis of MYC mRNA levels (relative to beta-actin) in DMEM treated control MiaPaCa2 cells, MiaPaCa2 cells treated with human recombinant FGF1 and, MiaPaCa2 cells treated with a combination of recombinant FGF1 and PD17307. **D,** qRT-PCR analysis of MYC mRNA levels (relative to beta-actin) in DMEM treated control MiaPaCa2 cells, MiaPaCa2 cells treated with CAF4414 derived CM, and MiaPaCa2 cells treated with a combination of CAF4414 derived CM and PD173074. $^{**}p < 0.01$ by one-way ANOVA ($n = 3$ biological replicates).

**Supplementary Figure 3: Paracrine FGF1 signaling augments MYC expression and stability via AKT/GSK-3β signaling.**

**A,** Western blots of whole cell and nuclear fractions isolated from MiaPaCa2 cells for the indicated signaling events after 3h treatment with DMEM or CAF 4422 derived CM. **B,** Western blots of whole cell and nuclear fractions isolated from 8988T cells for the indicated signaling events after 3h treatment with DMEM or CAF 4414 derived CM. **C,** Western blots for the indicated signaling events in MiaPaCa2 cells transfected with control plasmid or dominant-negative AKT and treated with DMEM or CAF 4414 CM for 3h. **D,** Western blots for the indicated MYC species in cells treated as in **C**. **E,** Western blots for the indicated signaling events in MiaPaCa2 cells transfected with control plasmid or constitutively active GSK-3β S9A and treated with DMEM or CAF 4414 CM for 3h. Western blots for the indicated MYC species were on nuclear lysates from cells under the same treatment conditions as for whole cell lysate analysis.

**Supplementary Figure 4: Stromal cues alter MYC chromatin occupancy and increase MYC target gene expression in PDAC cells.**

**A,** Gene ontology terms enriched among differentially bound loci by ChIP-seq (significantly enriched among CM-restricted binding sites). **B,** Heatmap showing hierarchical clustering of MYC binding sites identified under both DMEM and CM conditions by ChIP-seq as described in **A**. **C,** Results of HOMER analysis of top motifs among bound regions in MYC ChIP-seq (assessing all bound peaks, including DMEM and CM treatments). **D,** ChIP-qPCR for MYC binding sites in the indicated differentially bound promoters, as determined by ChIP-seq, in MiaPaCa2 cells treated with DMEM or CAF 4422 CM for 3h ($n = 2$ biological replicates). **E,** qRT-PCR for the indicated genes in MiaPaCa2 cells treated for 12h or 24h with CAF 4414 CM. Results were normalized to 36B4 as a housekeeping gene ($n = 3$ biological replicates). *$p < 0.05$, **$p < 0.01$, ***$p < 0.001$, ****$p < 0.0001$ by Student's t-test.

**Supplementary Figure 5: The FGF1/FGFR axis supports PDAC growth in vitro and MYC levels in vivo. A,** Proliferation assay on 8988T cells treated with CM from control or FGF1 knockdown CAF 4414 for 72h. **$p < 0.01$ by one-way ANOVA ($n = 3$ biological replicates). **B,** Proliferation assay for MiaPaCa2 cells treated with CM from control or FGF1 knockdown CAF 4414 for 72h. **$p < 0.01$ by one-way ANOVA ($n = 3$ biological replicates). **C,** Western blots for MYC and the indicated signaling events in PSN1 cells treated with CAF 4414 CM, BGJ 398 (2μM), and trametinib (20nM) for 3h.

Supplementary Figure 1

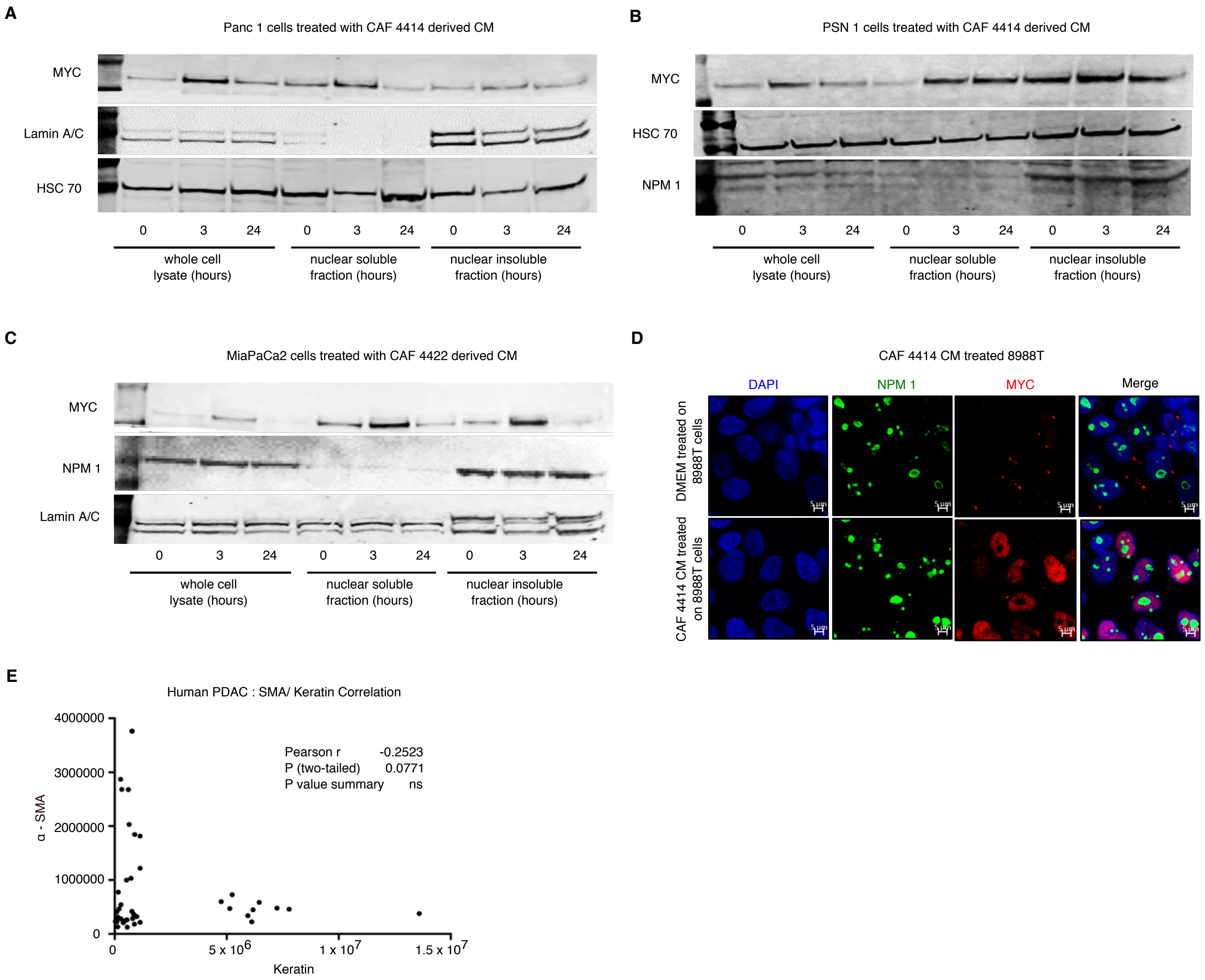

Supplementary Figure 2

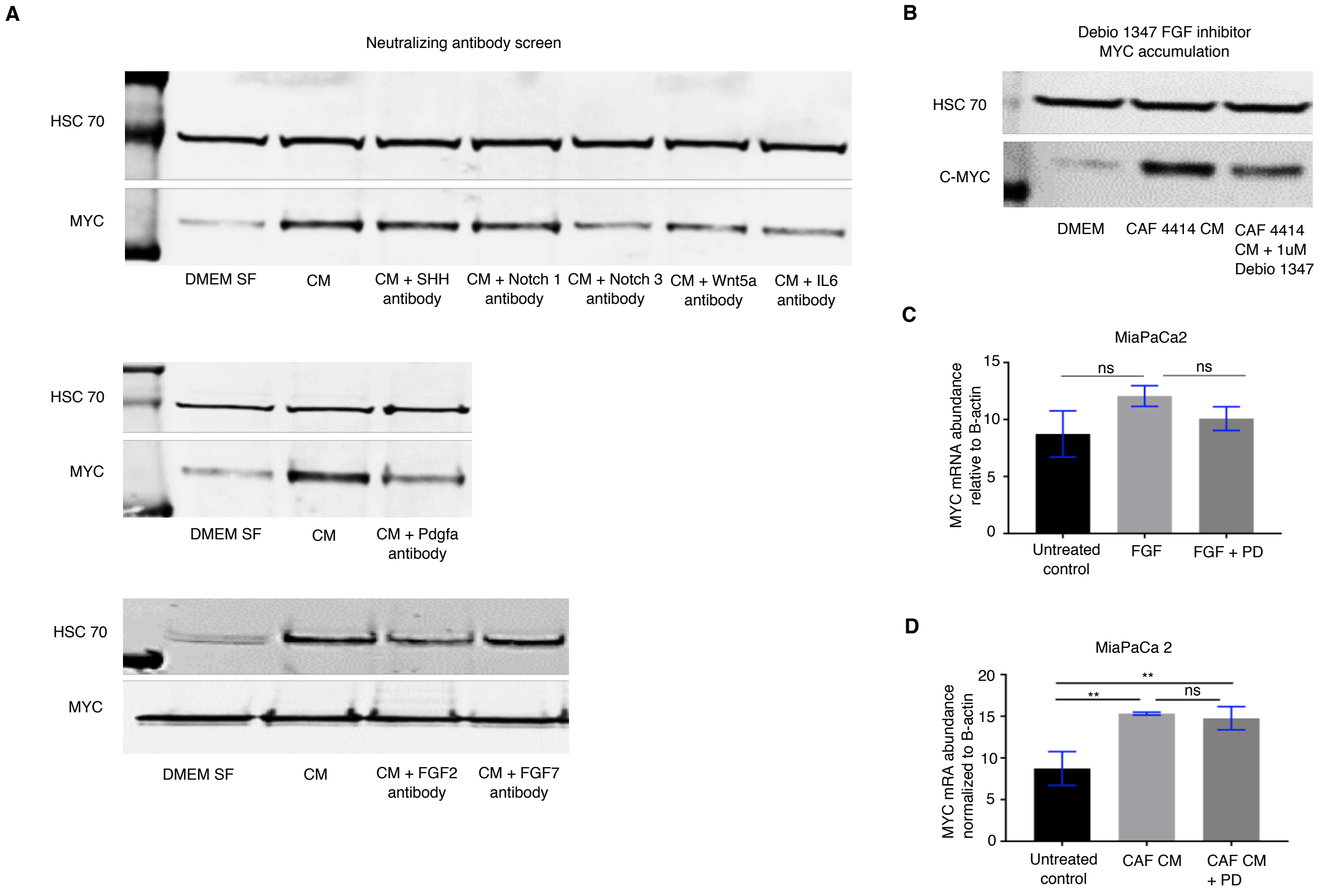

Supplementary Figure 3

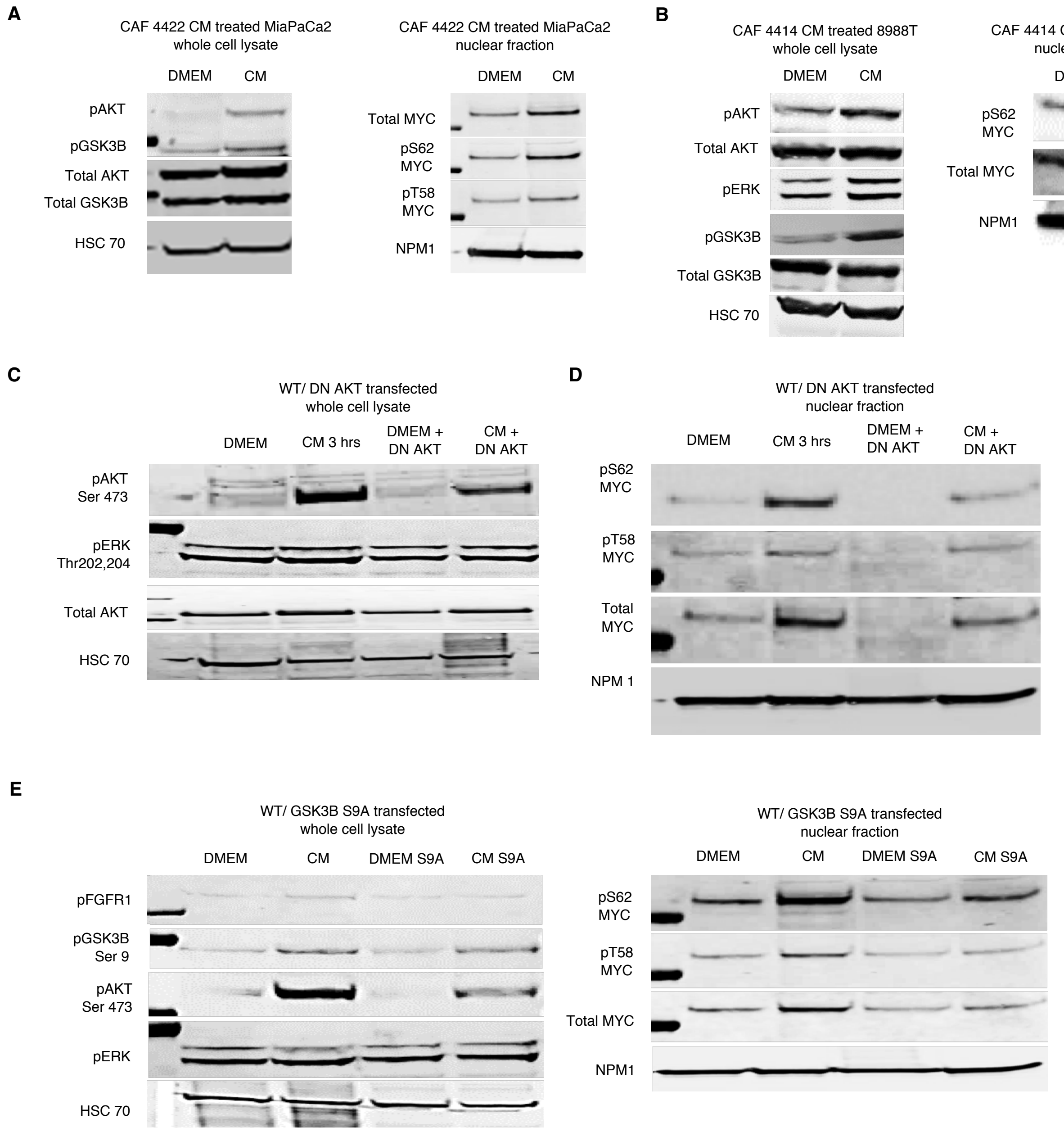

### Supplementary Figure 4

**A**

| GO term | p-value |
|---|---|
| ribosomal large subunit assembly | 1.8476E-05 |
| mitotic cell cycle | 0.00055245 |
| RNA splicing, via transesterification reactions | 0.00010324 |
| RNA splicing, via transesterification reactions with bulged adenosine as nucleophile | 0.00023633 |
| mRNA splicing, via spliceosome | 0.00023633 |
| RNA methylation | 0.00017918 |

**B**

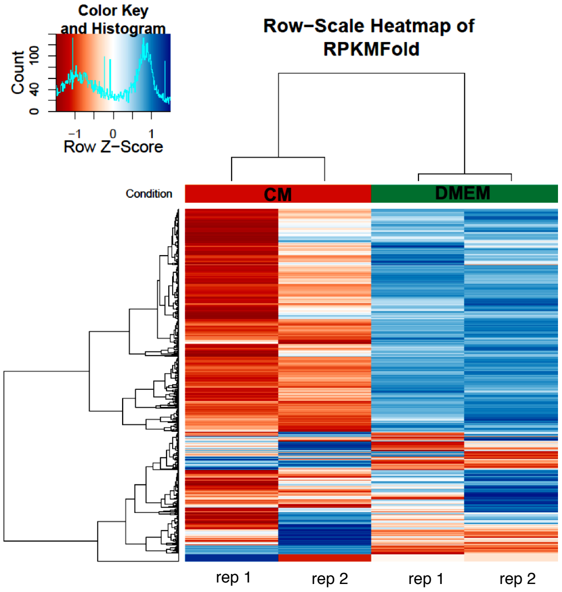


**C**

| Rank | Motif | P-value | log P-pvalue | % of Targets | % of Background | STD(Bg STD) | Best Match/Details | Motif File |
|---|---|---|---|---|---|---|---|---|
| 1 | 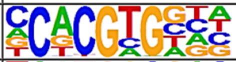 | 1e-100 | -2.303e+02 | 25.29% | 11.60% | 54.2bp (67.5bp) | c-Myc(bHLH)/LNCAP-cMyc-ChIP-Seq(Unpublished)/Homer(0.959) More Information \| Similar Motifs Found | motif file (matrix) |
| 2 | 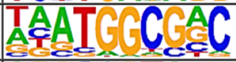 | 1e-28 | -6.627e+01 | 15.24% | 9.03% | 55.6bp (67.5bp) | YY2/MA0748.1/Jaspar(0.853) More Information \| Similar Motifs Found | motif file (matrix) |
| 3 | 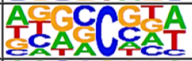 | 1e-27 | -6.431e+01 | 63.75% | 54.05% | 56.0bp (68.3bp) | PB0143.1_Klf7_2/Jaspar(0.616) More Information \| Similar Motifs Found | motif file (matrix) |
| 4 | 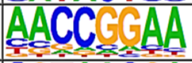 | 1e-23 | -5.511e+01 | 22.71% | 15.76% | 55.7bp (68.7bp) | ETV2/MA0762.1/Jaspar(0.914) More Information \| Similar Motifs Found | motif file (matrix) |
| 5 | 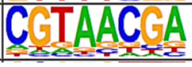 | 1e-22 | -5.089e+01 | 43.37% | 34.96% | 56.0bp (73.2bp) | PB0159.1_Rfx4_2/Jaspar(0.695) More Information \| Similar Motifs Found | motif file (matrix) |

**D**

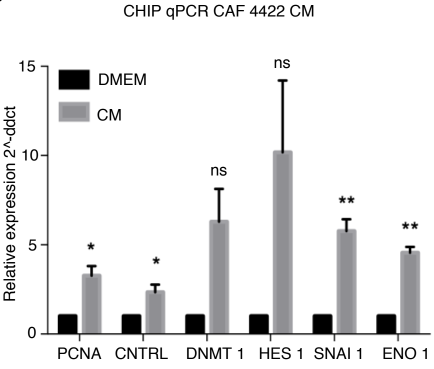


**E**

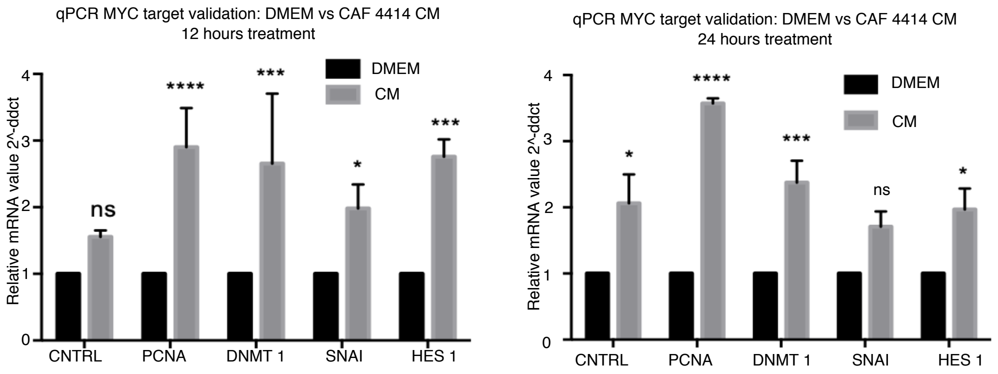

Supplementary Figure 5

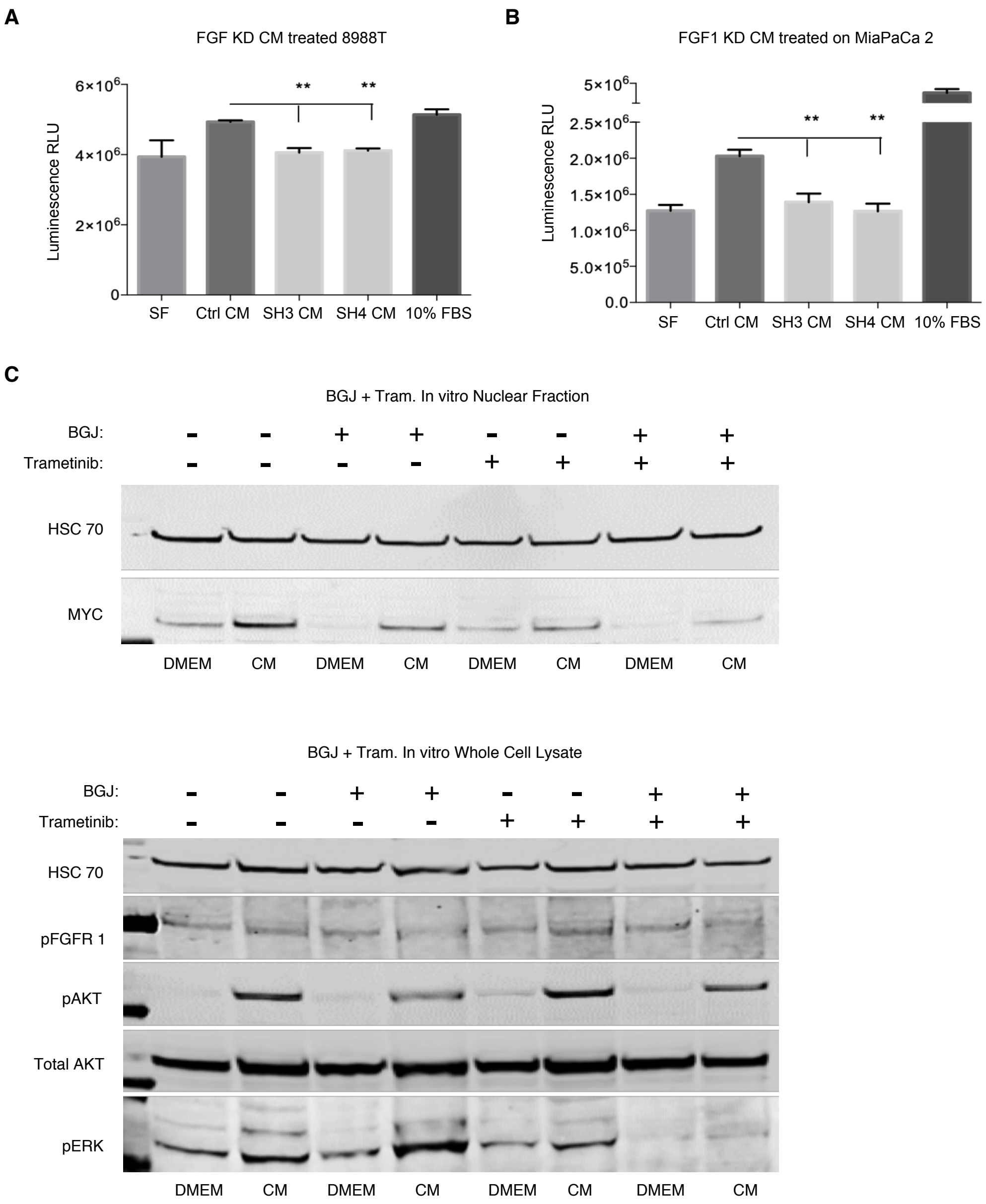

Acidic fibroblast growth factor underlies microenvironmental regulation of MYC in pancreatic cancer

#### Supplemental Materials and Methods

##### Generation of human PDAC primary Cancer Associated Fibroblasts (CAFs)

All of the primary fibroblast strains were derived in the same way from pancreatic tumor explant cultures. Tissue samples from primary pancreatic tumor were cut into small pieces approximately 2-3 $mm^3$ in size and seeded in fetal calf serum medium (FCS) with 200u/ml Penicillin, 200ug/ml Streptomycin, 100ug/ml Gentamicin and 2.5g/ml Amphotericin B. When outgrowths of fibroblasts appeared, the culture medium was replaced with complete F medium to facilitate fibroblast growth. The remnants of the tissue were carefully washed away and CAFs were routinely maintained in complete F medium at 37ºC in a humid atmosphere containing 5% CO2. Immunohistochemistry and RNA-seq were used to confirm minimal expression of cytokeratin and abundant expression of $\alpha$SMA in primary CAF lines. Wild-type *KRAS* codon 12 sequence was confirmed for all CAF lines and mutant *KRAS*-G12 was confirmed in all corresponding parent tumors. All experiments in this study were performed with CAFs at 7 passages at most.

##### Pancreatic cancer cell lines

Human pancreatic cancer cell lines MIAPaCa-1, PA-TU-8988T and PSN-1 were obtained from ATCC and grown in Dulbecco's modified Eagle's medium (DMEM) containing 10% fetal bovine serum.

**Western Blotting**

PDAC cells were serum starved for 48 hours and treated as mentioned in the text. Sub-cellular fractions were prepared using the NE-PER® Nuclear and Cytoplasmic Extraction reagents (Thermo Scientific) as per manufacturer's instruction. Protein concentration was quantitated using the BCA protein assay kit (Pierce). Equal amounts of protein were loaded in each lane and separated on a 4-12% Bis-Tris NuPAGE® gel (Invitrogen), then transferred onto a PVDF membrane. Membranes were probed with primary antibodies and infrared secondary antibodies: IRDye 700 goat anti-rabbit IgG or IRDye 800 goat anti-mouse IgG (LI-COR Biosciences). For protein band quantitation, infrared signals were detected using the Odyssey CLx infrared imaging system and bands quantified using Image Studio software (LICOR Biosciences).

**Neutralizing antibody screen**

Specific neutralizing antibodies were added to CAF derived CM and rocked at room temperature for 1 hour for effective neutralization of CAF secreted factors. PDAC target cells were serum starved for 48 hours and then treated with CAF derived CM only or CAF derived CM with specific neutralizing antibodies. The complete list of neutralizing antibodies used is described under the antibodies section of Materials and Methods.

**Immunostaining**

Cells: Cells plated on coverslips were fixed in 4% paraformaldehyde for 15 minutes at room temperature, washed three times with PBS, and permeabilized with .1% Triton X-100 for 10 min at room temperature. Following permeabilization, coverslips were blocked for one hour at room temperature in blocking solution (8% BSA solution) and then transferred to a carrier solution (8% BSA solution) containing diluted antibodies (described in antibody section of Supplementary Materials and Methods). Coverslips were incubated with the primary antibody for 3 hours at room temperature and then washed five times for 5 minutes each in PBS following which, secondary Alexa-flour conjugated antibodies diluted in the same carrier solution (1:400) were added to the coverslips for one hour at room temperature. After the secondary antibody incubation, coverslips were washed five times for five minutes each in PBS and mounted with Vectashield mounting media containing DAPI.

Mouse and Human tissue sections: Mice were anesthetized and euthanized according to institutional guidelines. Pancreatic tumors/subcutaneous tumors were excised carefully and drop fixed overnight in 4% paraformaldehyde (PFA). Tissue samples were paraffin embedded, sectioned and H&E stained at OHSU Histopathology Core. In brief, tissue sections were de-paraffinized and rehydrated through an ethanol series and ultimately in PBS. Following antigen retrieval, tissue samples were blocked for two hours at room temperature in blocking solution (Aqua Block buffer from Abcam) and then transferred to a carrier solution (Aqua Block buffer from Abcam) containing diluted antibodies (described in antibody section of Supplementary Materials and Methods). Sections were incubated overnight at room temperature and then washed five times for 5 minutes each in PBS following which, secondary Alexa-flour conjugated antibodies diluted in the same carrier solution (1:400) were added to the sections for one hour at room temperature. Sections were then washed five times for five minutes each in PBS and were mounted with Vectashield mounting media containing DAPI. To stain for MYC, human PDAC sections were stained with a MYC pS62 antibody (Abcam #ab185656) while subcutaneous and orthotopic tumors in mice were stained with a total MYC antibody (CST #5605). Additional antibodies are listed in the Antibodies section.

**2-plex Fluorescence In Situ Hybridization**

Human PDAC tumor tissue samples were subjected to Fluorescence In Situ Hybridization using the ViewRNA™ ISH Tissue Assay Kit (2-plex) from Thermo Fisher Scientific according to manufacturer guidelines. In brief, samples were subjected to controlled protease digestion for initial permeabilization following which, the samples were incubated with the proprietary probe-containing solution according to the manufacturer guidelines. It was crucial that samples remained fully submerged during the entire incubation. Following probe hybridization, samples were washed and then subjected to sequential hybridization with pre-amplifier and amplifier DNA and fluorophore. Hybridizations with pre-amplifier, amplifier, and fluorophore were performed as indicated by the manufacturer. Samples were mounted using DAKO ultra mount mounting medium as indicated by the manufacturer.

**Microscopy**

Imaging of fluorescence staining was done by confocal imaging of fixed cells and tissues with a laser-scanning confocal inverted microscope (LSM 880, Carl Zeiss, Inc.) and a 40×/ 1.1 NA water objective or 63x/1.4 NA oil objective was used to image the samples. A Zeiss Axio Scan automated slide scanning microscope was used to take 20x tiled images of entire H&E stained mouse tumor sections.

**ChIP-seq**

Immunoprecipitation and sequencing: Chromatin immunoprecipitation was performed as described previously (1). Briefly, PDAC cells were fixed in 1% formaldehyde, and nuclei were isolated and lysed in buffer containing 1% SDS, 10 mM EDTA, 50 mM Tris-HCl pH 8.0, and protease inhibitors, and sheared with a Diagenode Bioruptor to chromatin fragment sizes of 200 – 1000 base pairs. Chromatin was immunoprecipitated with antibodies to MYC (CST # 9402), or rabbit IgG (Santa Cruz Biotechnology). ChIP-seq libraries were constructed using Illumina's TruSeq ChIP library preparation protocol, using 5ng input DNA. Libraries were sequenced using 100 cycle single read sequencing on a HiSeq 2500.

Alignment and Peak Calling: Reads were first trimmed of adapter sequences using default parameters of Trim Galore! (version 0.4.3) (retrieved from https://www.bioinformatics.babraham.ac.uk/projects/trim_galore/). Adapter-trimmed reads were aligned to the human reference genome (GRCh38, release 87) using bowtie2 (version 2.3.4) (2). The resulting alignment files were further filtered to contain reads with a MAPQ score of 11 or greater using samtools (version 1.3.1) (3) and all duplicates were removed using picard tools (version 2.9.0) (http://broadinstitute.github.io/picard). Peaks were called using the "callpeak" method of MACS2 (version 2.1.1.20160309) (4) with a q-value cut-off of 0.05.

Overlapping Peaks: Peaks output by MACS2 were filtered for known chromosomes only. The findOverlapsOfPeaks() function from the ChIPpeakAnno R package (version 3.14.2) (5) was used to determine peaks shared between replicates within a treatment. The within-treatment consensus peaks were similarly evaluated in order to create a final set of peaks shared between both treatments.

The resulting between-treatment consensus peaks were evaluated for motif enrichment using the findMotifsGenome.pl script from the homer package (version 4.10.1) (6). The read counts of the ChIP reads relative to the control reads were used to visualize the binding density of the between-treatment consensus peaks using the dba.plotHeatmap() function from the diffBind R package (version 2.8.0) (https://bioconductor.org/packages/release/bioc/html/DiffBind.html).

Gain of MYC binding at known MYC target gene promoter sequences was confirmed by ChIP qPCR. Enrichment was calculated against DMEM treated samples and enrichment values were normalized to a control intergenic region of the genome. The primer sequences used are described in the sequences section of Online Methods.

**Gene Expression Analysis by qPCR**

The isolated total RNA (1 $\mu$g) was reverse-transcribed to produce cDNA using iScript Reverse Transcription Supermix kit (Bio-Rad, Hercules, CA, USA). Real-time PCR was performed using SYBR Green supermix (Bio-Rad). The cDNA sequences for specific gene targets were obtained from the human genome assembly (http://genome.ucsc.edu) and gene specific primer pairs were designed using the Primer3 program (http://frodo.wi.mit.edu/primer3/primer3_code.html). Relative gene expression was expressed as fold change in gene expression between non-tumor pancreatic stellate cells and primary pancreatic tumor associated CAFs or DMEM treated cells and CM treated cells, with threshold cycle (CT) values normalized using the 36B4 housekeeping gene. Gene specific primer pair sequences are provided under primer section of Supplementary Materials and Methods.

**ELISA**

Secreted human acidic FGF in CM was measured using a sandwich immune-luminometric assay based on manufacturer instructions (Human FGF acidic Quantikine Elisa Kit, R&D # DFA00B). The concentration of secreted acidic FGF was compared across a subset of primary pancreatic tumor derived CAF strains and a panel of PDAC cell lines.

**Stable and Transient Plasmid transfections**

The vectors containing shRNA against human acidic FGF were purchased from Dharmacon. In brief, fifteen micrograms of vector, together with 7.5 $\mu$g of each packaging vector (pMD2.G and psPAX2), were co-transfected into 293T cells. Supernatant-containing lentivirus particles were harvested 48 hours and 72 hours after transfection, passed through a 0.45-μm membrane filter, and directly used to infect primary pancreatic tumor associated CAFS in the presence of 6 μg/mL polybrene. After 48 hours, the infected cells were maintained in the medium with 2 μg/mL puromycin for 2 weeks before knockdown assessment.

For transient transfections with purified plasmid DNA, transfections were performed using Lipofectamine 2000 following the manufacturer's instruction.

**Proliferation assay**

For the growth assays, PDAC cells were seeded into 96-well plates at $2x10^3$ cells per well in DMEM containing 10% FBS. The next day, cells were washed with PBS and changed to serum free DMEM containing 25 mM glucose and 4 mM glutamine for 72 hours. Post serum starvation, cells were treated with CM -/+ pharmacological inhibitors (as mentioned) at the indicated concentrations for another 72 hours. After 72 hours, cells were lysed with CellTiter-Glo® Luminescent Cell Viability Assay reagent (Promega) and luminescence was read using the GloMax plate reader.

**Antibodies**

Immunofluorescence studies

Smooth Muscle Actin Monoclonal Antibody [1A4 (asm-1)] – Invitrogen # MA5-11547

Anti c-Myc antibody [Y69] – Abcam # ab32072

Anti c-Myc (pS62) antibody [EPR17924] – Abcam # ab185656

Anti c-Myc (D84C12) Rabbit mAb – CST # 5605

Anti-FGFR1 (phospho Y653) antibody [EPR843 (N)] – Abcam #ab173305

FGFR1 Monoclonal Antibody (VBS-7) – Invitrogen # 13-3100

NPM1 Monoclonal Antibody (FC-61991) – Invitrogen # 32-5200

Pan Keratin (C11) Mouse mAb (Alexa Fluor® 647 Conjugate) – CST#4528

Cytokeratin Pan Type I/II Antibody Cocktail – Thermo Fisher Scientific # MA5-13156

Western Blots

c-Myc (D84C12) Rabbit mAb – CST #5605

Anti c-Myc (pS62) antibody [EPR17924] – Abcam # ab185656

Anti c-Myc (pT58) antibody [EPR17924] – Abcam # ab28842

Phospho-Akt (Ser473) (D9E) XP® Rabbit mAb – CST #4060

Akt (pan) (C67E7) Rabbit mAb – CST #4691

Phospho-GSK-3β (Ser9) Antibody – CST #9336

GSK-3β (27C10) Rabbit mAb – CST #9315

Lamin A/C (4C11) Mouse mAb – CST #4777

HSC 70 mouse Antibody (B-6) – SC #7298

NPM1 Monoclonal Antibody (FC-61991) – Invitrogen # 32-5200

Human FGF acidic neutralizing Antibody – R&D # AF232

Human FGF basic neutralizing Antibody – R&D # AF232

Human FGF-7 neutralizing Antibody – R&D # MAB251

Human Il-6 neutralizing Antibody – R&D # MAB2061

Human HGF neutralizing Antibody – R&D # AB-294-NA

Human PDGF-AA neutralizing Antibody – R&D # MAB221

Human PDGF-BB neutralizing Antibody – R&D # AB-220-NA

Human/Mouse Wnt-3a neutralizing Antibody – R&D # MAB9025

**Plasmids**

HA-AKT DN (K17M) was a gift from Mien-Chie Hung (Addgene plasmid#16243).

HA-GSK3 beta S9A pcDNA3 was a gift from Jim Woodgett (Addgene plasmid #14754).

SMARTvector Lentiviral Human shRNA constructs were purchased from Dharmacon (#224940601, #225482233, #226573540, #228128665). Flag-tagged MYC WT and T58A plasmids were kindly provided by Dr. Mushui Dai (OHSU).

**Primer Sequences**

Gene Expression analyses

Human MYC F: CAGCTGCTTAGACGCTGGATT; R: GTAGAAATACGGCTGCACCGA

Human FGF1 F: CAATGTTTGGGCTAAGACCTG; R: GGCTGTGAAGGTGGTGATTT

Human FGF2 F: GTGTGTGCTAACCGTTACCT; R: GCTCTTAGCAGACATTGGAAG

Human FGF7F: ATCAGGACAGTGGCAGTTGGA; R: AACATTTCCCCTCCGTTGTGT

Human CSF1 F: GCTCTCCCAGGATCTCATCAC; R: TCAAAGGAACGGAGTTAAAACGG

Human EGF F: CTGTGGTGCTGTCATCTGTC; R: GTCCCCAGCCGATTCCTTG

Human HGF F: AAGGTGACTCTGAATGAGTC; R: GGCACATCCACGACCAGGAACAATG

Human NGF F: CACACTGAGGTGCATAGCGT; R: TGATGACCGCTTGCTCCTGT

Human PDGFAA F: CACACCTCCTCGCTG TAGTATTTA; R: GTTATCGGTGTAAATGTCATCCAA

Human PDGFBB F: TCCCGAGGAGCTTTATGAGA; R: ACTGCACGTTGCGGTTGT

Human KITLG F: CAGAGTCAG TGTCACAAAACCATT; R: TTGGCCTTCCTATTACTGCTACTG

Human TGFB 1 F: GCAACAATTCCTGGCGATACCTC; R: AGTTCTTCTCCGTGGAGCTGAAG

Human TGFB 2 F: AGAGTGCCTGAACAACGGATT; R: CCATTCGCCTTCTGCTCTT

Human VEGFA F: GCCTTGCTGCTCTACCTCCA; R: CAAGGCCCACAGGGATTTT

Human VEGFB F: AGCACCAAGTCCGGATG; R: GTCTGGCTTCACAGCACTG

Human IL-6 F: AAAGAGGCACTGGCAGAAAA; R: AGCTCTGGCTTGTTCCTCAC

Human IL-8 F: CTGGCCGTGGCTCTCTTG; R: CCTTGGCAAAACTGCACCTT

Human IL-1a F: AGGGAATTCACCCCAAGAAC; R: ACTATGGGGGATGCAGGATT

Human IL-1b F: AAGCTGAGGAAGATGCTG; R: ATCTACACTCTCCAGCTG

Human IL-32 F: ATGTGCTTCCCGAAGGTCCTCTCTGA; R: TCATTTTGAGGATTGGGGTTCAGAGC

Human CXCL1 F: AGGGAATTCACCCCAAGAAC; R: ACTATGGGGGATGCAGGATT

Human WNT5a F: AGAAGAAACTGTGCCACTTGTATCAG; R: CCTTCGATGTCGGAATTGATACT

Human MIF F: CGCAGAACCGCTCCTACAG; R: GGAGTTGTTCCAGCCCACAT

Human beta actin F: ATTGGCAATGAGCGGTTCCGC; R: CTCCTGCTTGCTGATCCACATC

Human 36B4 F: GTGCTGATGGGCAAGAAC; R: AGGTCCTCCTTGGTGAAC

MYC target validation

Human CNTRL F: CAGATGAAAGCCCTTACATTGGC; R: CTGCCTGAGCACTGTCAATAAT

Human PCNA F: AGGGCTCCATCCTCAAGAAG; R: GTAGGTGTCGAAGCCCTCAG

Human DNMT1 F: TCAGGGACCACATCTGTAAGG; R: GCCGTTCTTCCTGTCATGG

Human SNAI1 F: CTAGAGTCTGAGATGCCCCG; R: AGTTCTGGGAGACACATCGG

Human EIF2B5 F: AGAGGCGAACTTCACTGACA; R: GTCCAGCACCACGTTATCAC

Human HES1 F: GCTTTCCTCATTCCCAACGG; R: GTGGGTTGGGGAGTTTAGGA

ChIP qPCR validation

Human DNMT1 F: TGCAATGAATTCCAGATGTG; R: GGAGGGGCAGAGTGAGAGAT

Human PCNA F: TTGGCTAATCGCACACTGA; R: GTCCGGAATATCCACCAATG

Human SNAI1 F: GGGCCTTTTCCCTTGATAAT; R: AAGGGAAGTGTGCTTTGGTG

Human HES1 F: AAGTTTCACACGAGCCGTTC; R: GAGAGGTAGACGGGGGATC

Human CNTRL F: CCTTCCAACAGTACCGGAGA; R: GCAGCCATTTTGTTGTGTTG

**Pharmacological compounds**

Cycloheximide was purchased from Cell Signaling Technologies: CST #2112

Actinomycin D was purchased from Sigma-Aldrich: # A1410-5MG

PD173074 was purchased from Selleck Chemicals: # S1264

NVP-BGJ398 was purchased from Selleck Chemicals: # S2183

MK-2206 2HCl was purchased from Selleck Chemicals: # S1078

Trametinib was purchased from MedChem Express: # HY-10999A

Purified Recombinant Human FGF acidic was purchased from Stem Cell Technologies: # 78187.1

All pharmacological compounds were resuspended and stored according to manufacturer guidelines.

Complete F media for growing and maintenance of primary human PDAC associated CAFs:

3:1 DMEM/F12

5% FBS

0.4 ug/ml hydrocortisone

5 ug/ml insulin

8.4 ng/ml cholera toxin

10 ng/ml EGF

24 ug/ml Adenine

10 uM ROCK inhibitor
